# A female heterogametic ZW sex-determination system in Acariformes

**DOI:** 10.1101/2023.10.24.563255

**Authors:** Svenja Wulsch, Hüsna Öztoprak, Nadège Guiglielmoni, Daniel L. Jeffries, Jens Bast

## Abstract

Sexual reproduction, while often associated with separate sexes, is an ancient and widespread feature of multicellular eukaryotes. While a diversity of sex-determination mechanisms exist, for many organisms, which of these mechanisms is used remains unknown. Exploring sex-determination mechanisms in Acariformes, among the oldest chelicerate clades, is intriguing due to its potential to unveil conserved sex-determination systems. This insight can have implications for understanding sex chromosome evolution and its broader impact on higher taxa.To identify the mechanism of sex determination in Acari, i.e., oribatid mites, we generated a high-quality chromosome-level genome assembly of *Hermannia gibba* (Koch, 1839) by combining PacBio HiFi and Hi-C sequencing. Coverage and allele-frequency analyses on pools of male and female individuals suggest a female-heterogametic ZW sex-determination system with little degeneration of the W chromosome. To date, this represents the only documented case of a ZW system in Acariformes. Further comparative studies in H. *gibba* will reveal how old the ZW system is and whether it exhibits conservation or polymorphism.

## Introduction

The vast majority of metazoans reproduce sexually whereby two separate sexes mix their genomes via meiosis and fusion of gametes to generate offspring (Bell 1982). Despite this being such a widely shared and fundamental feature, the mechanisms that determine whether individuals develop into males or females are highly diverse (Bachtrog et al. 2014; Beukeboom and Perrin 2014). Two (not mutually exclusive) main types of sex determination occur in nature: environmental sex determination (ESD), where environmental cues trigger sex development; and genotypic sex determination (GSD), where sex-specific genotypes determine the individual’s sex, such as sex chromosomes. Sex chromosome systems have evolved independently numerous times throughout eukaryotes. The spectrum ranges from stable sex chromosomes, e.g. in birds or mammals (Bull 1984; Zhou et al. 2014) to rapidly evolving sex chromosome pairs with frequent turnovers, e.g. in frogs and fish (Takehana et al. 2014; Jeffries et al. 2018). Why the stability of sex chromosomes and the prevalence of transitions differ so much across the tree of life remains one of the unsolved mysteries in the field of reproductive systems evolution. To answer this question, it is imperative to characterize the diversity of sex chromosome systems across a diversity of organisms (Bachtrog et al. 2014; Consortium et al. 2014; Palmer et al. 2019).

Acariformes are one of the major groups of Acari including over 11,500 species with many more undescribed (Subías 2023). They likely originated during the Precambrian and were among the first to colonize land (Schaefer et al. 2010; Lozano-Fernandez et al. 2019). Despite their age and diversity, relatively little is known of their sex-determination system. Generally, there is considerable debate and contrasting reports about sex determination in Acariformes based on karyotypes (Heethoff et al. 2006). The ancestral state in Acariformes is believed to be diplodiploidy, i.e. the same karyotype for females and males without distinct sex chromosomes (Sokolov 1954; Norton et al. 1993; Wrensch et al. 1994). Most previous studies on the major Acari groups (Parasitiformes and Acarifomres) investigated sex determination based on indirect evidence or karyotypes: in Parasitiformes 76 cases with XO or XY sex chromosomes and 106 with some form of haplodiploid, whereas in Acariformes 17 cases of XO or XY are reported and 106 with a type of haplodiploidy (Heethoff et al. 2006; Consortium et al. 2014).

Oribatid mites (Acariformes, Acari), lack differentiated sex chromosomes and are diploid (mostly 2n=18) (Norton et al. 1993). Due to conflicting evidence hinting at the potential existence of haplodiploidy or parahaploidy, the exact sex-determination mechanism present in oribatid mites remains elusive (Oliver 1983; Norton et al. 1993; Wrensch et al. 1994). The sex ratios of some sexual oribatid mites are balanced (1:1) or they tend to show a female bias (Grandjean 1941; Webb and Gw 1979; Smelansky 2006; Wehner et al. 2014). Moreover, the presence of diverse sex-determination systems within the group cannot be ruled out, given its large size, diversity, and the paucity of prior investigations into sex-determination mechanisms. In most species where balanced sex ratios are observed, sex is genetically determined through the mechanism of GSD and often involves sex chromosomes (Uller et al. 2007). Some oribatid mite species show an equal 1:1 sex ratio, whereas for most species female-biased (72%-69%) sex ratios are observed (Domes et al. 2007). Interestingly, a large number of asexual species exist in the Oribatida that reproduce in the absence of sex over a substantial amount of time (Brandt et al. 2021; Wehner et al. 2021; Öztoprak et al. 2023).

*Hermannia gibba* (Hermanniidae, Sarcoptiformes), a widely distributed sexual species found in diverse forest soils, is presumed to employ genetic sex determination due to the unstable nature of its soil habitat with various microhabitats (Smrž 2010; Liana and Witaliński 2012; Lóšková et al. 2013). However, a comprehensive understanding of its sex-determination mechanisms remains elusive. In this study, we investigate the sex-determination system of the oribatid mite *H. gibba*. Prerequisites for the identification of the sex-determining region (SDR) are high-quality genome and sequence data of pools from male and female individuals.

Thus, we first generated a chromosome-level genome assembly from a female *H. gibba* individual up to current standards (Guiglielmoni et. al 2022). Second, we conducted whole-genome sequencing of pools of female and male individuals. We used two complementary approaches to identify regions of sex linkage (Palmer et al. 2019): scans of i) coverage differences and ii) allele frequencies along the genome of male and female mite pools. Further for additional verification, we performed a restriction enzyme digestion of a predicted female-specific site and confirmed the adequacy between morphological and molecular identifications (Methods and Figure S4). Our results identify a ZW female-heterogametic sex-determination system with a sex-linked region of 14.54 Mb on Chromosome 1 in *H. gibba*. To date, this is the first documented case of a ZW system in Acari.

## Results

### Genome assembly and annotation

The *de novo* genome assembly of *Hermannia gibba* was performed using a combination of Pacific Biosciences (PacBio) HiFi reads from a single female individual, and Hi-C reads from a population pool, as described in **Figure 1a** (see Methods for details). PacBio HiFi reads were first analyzed with *k*-mer spectra, which showed that sequencing data had little contamination and the genome was predicted as diploid (Supplementary Figures S1, S2). The initial assembly resulted in 934 contigs with a total size of 537.1 Mb, which was decreased to 415 contigs and a total size of 371.1 Mb after haplotig purging. Subsequent Hi-C scaffolding yielded eight chromosome-level scaffolds. The final assembly has a size of 352.2 Mb, 61 gaps, and with few duplicated regions (**Figure 1b**). The Benchmarking Universal Single-Copy Orthologs (BUSCO) tool was used to assess completeness, resulting in a score of 96.1% against the Arachnida odb10 database (92.7% single-copy orthologs, 3.4% duplicated orthologs). The eight chromosome candidates, ranging from 29.5 to 75.9 Mb in size, can be identified in the Hi-C contact map (**Figure 1c**). 55.92% of the assembly was identified as repetitive content, including 29,072 telomeric repeats of the sequence ‘AACCT’. Genome annotation yielded 25,869 predicted protein-coding genes with a BUSCO score of 95.5% against the Arachnida odb10 database (91.5% single-copy orthologs, 4.0% duplicated orthologs).

**Figure 1.**
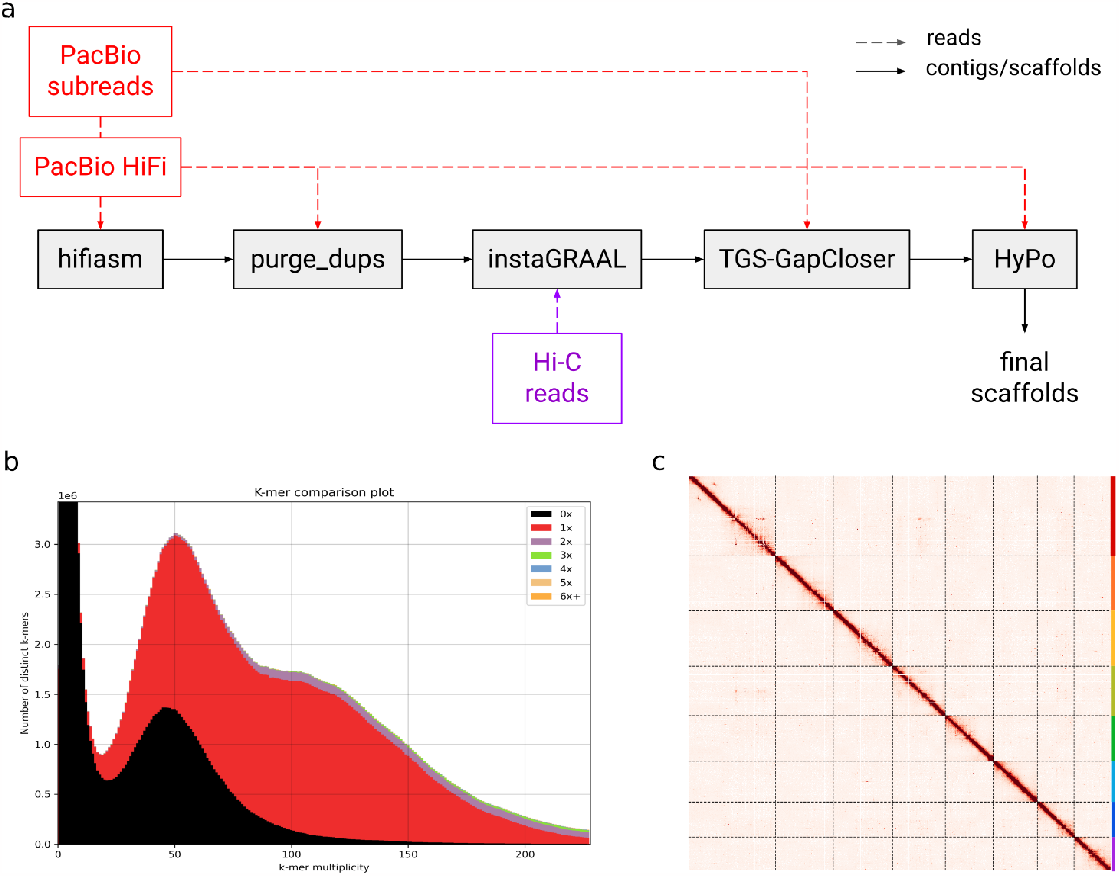
Genome assembly of female *Hermannia gibba*. a) Assembly workflow. b) *k*-mer analysis of the final assembly. Low-multiplicity *k*-mers are absent from the assembly (0X). The heterozygous peak and the homozygous peak are respectively identified around 50X and 110X. As haplotypes are collapsed in the assembly, homozygous *k*-mers are represented once and only half of heterozygous *k*-mers are included in the assembly. There are limited artifactual duplications (2X in the homozygous peak). c) Hi-C contact map of the eight chromosome candidates.

### Searching for degenerated sex-linked regions using read coverage

To identify the SDR, we searched for genomic differences between females and males by sequencing pools of 30 individuals of each sex from *H. gibba* (for details see methods, Figure S1).

Sex chromosomes often lose recombination between gametologs (i.e. X/Y or Z/W) in the region surrounding the sex determination locus. These non-recombining regions are expected to accumulate sequence divergence over time, as well as deleterious mutations, and repetitive sequences. Given enough time, the sex-limited chromosome (Y or W) eventually loses large numbers of genes (Charlesworth et al. 2005; Jeffries et al. 2018), resulting in regions of the X or Z that are haploid in the heterogametic sex. To search for such highly diverged and/or degenerated sex-linked regions, we aligned the pooled sequencing data against the female repeat-masked reference genome, yielding comparable mapping rates of the female pool (96.76%) and the male pool (98.11%). We then compared coverage between male and female pools in 1 kb windows along the genome. If degenerated sex-linked regions exist, we would expect to find a male/female coverage ratio Δ = 0.5 for a XY system, or Δ = 2 for a ZW system. While autosomal, pseudoautosomal, or undegenerated sex-linked regions are expected to have a male/female coverage ratio of Δ = 1. The ratio of male-to-female coverage along the genome of *H. gibba* did not show any differences, indicating either a lack of sex chromosomes or the absence of a degenerated sex-linked region (**Figure 2a**).

**Figure 2.**
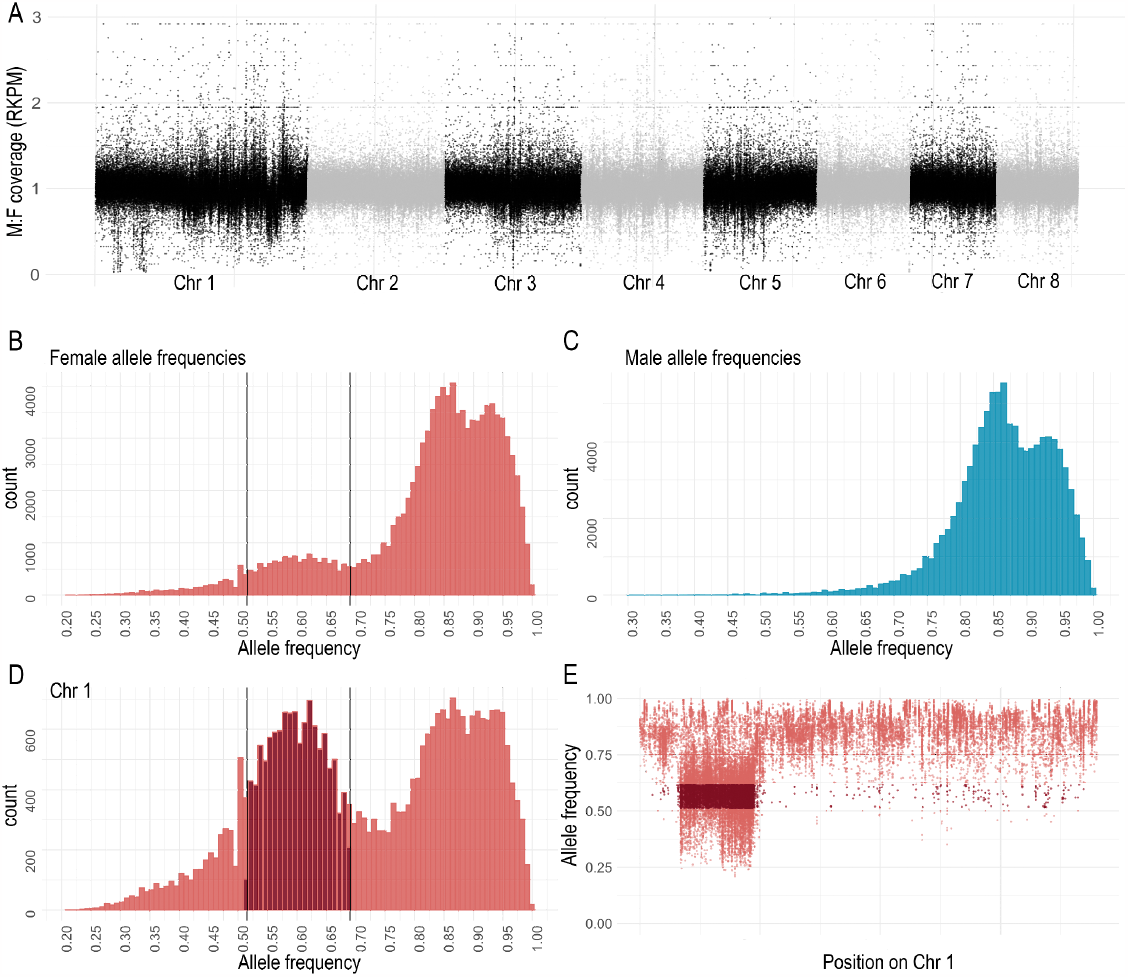
The sex determination region is not highly degenerated in *Hermannia gibba*. RPKM normalized coverage differences along the eight chromosomes in 1 kb blocks do not reveal any significant differences in coverage between male and female genomes (A). The histograms (B) and (C) provide an overview of the allele frequency distributions within each sex across the genome. The x-axis represents the allele frequency, and the y-axis represents the number of alleles corresponding to each frequency. Only alleles with an allele frequency (AF) greater than 0.95 in the other sex are included. The chromosome-specific histogram (E) shows the allele frequencies on Chromosome 1, with allele frequencies falling within the calculated threshold of 0.51 to 0.69 displayed in dark red. Scatter plot (E) visualizes the position of allele frequencies falling within the calculated threshold range on Chromosome 1. These allele frequencies indicate a sex-linked region spanning a continuous region of 14.54 Mb.

### Identification of sex-linked regions using allele frequency calculation

Nucleotide sequences in SDRs can diverge more rapidly between the gametologs than large changes in chromosome structure. Thus, in sex-linked regions that recently lost recombination and have not yet degenerated to the point of exhibiting coverage differences, differences in allele frequencies (AF) between males and females can be more informative. We found 4,663,024 polymorphic sites between the two pools which we used to search for loci with SNPs specific to the sex-limited chromosome. Such loci are expected to possess a major allele frequency of 1 in the homogametic sex, and 0.5 in the heterogametic sex. We therefore filtered for loci where one of the sexes is fixed (with a major allele frequency of > 0.95) and the other to be half (∼0.5; **Figure 2b**,**c**,**d)**. As we did not know the system of heterogamety before performing this analysis, we did this twice, once to search for an XY system by filtering for a major allele frequency of >0.95 in females and ∼0.5 in males, and once to test for a ZW system with a major allele frequency of >0.95 in males, and ∼0.5 in females.

Only in the female pool, we observed loci that exhibited allele frequencies ranging from 0.51 to 0.69, calculated based on the local maxima and minima. Among these 7,649 discovered SNPs, 7,029 (92%) were located on Chromosome 1, forming a contiguous region of 14.54 Mb from the start of 6.3 Mb to the end of 20.8 Mb (**Figure 2e**). As these loci are homozygous in males and heterozygous in females, we inferred the presence of a ZW female heterogametic genotypic sex-determination system. To test this hypothesis, we searched for the presence of a sex-specific locus in the sex-linked region that is exclusively present in females and absent in males. We performed restriction enzyme digestion, specifically targeting the female-specific locus located within a HindIII restriction site (Figure S4).

## Discussion

Historically, sex determination in oribatid mites relied mainly on cytological approaches, which led to contradicting results (Heethoff et al. 2006). Our study represents the first time a sequencing-based approach has been used to identify the sex-determination system within the Acariformes group, while also uncovering the first documented case of a ZW system in Acariformes. Our investigation revealed a lack of extensive degeneration of the *Hermannia gibba* sex chromosome. This may reflect a relatively recent loss of recombination in the sex-linked region. However, the age of the sex chromosome remains unknown as there is a large variation in the rate at which sex chromosomes lose recombination, and in the rate at which they degenerate once recombination has ceased (Wright et al. 2016).

The relatively large size (75.9 Mb) of the identified sex chromosome compared to autosomes might indicate chromosomal rearrangements. Oribatid mites possess holocentric chromosomes, which, unlike monocentric chromosomes, offer unique advantages that may impact recombination patterns (Wrensch et al 1994, Izraylevich 1995). Holocentric chromosomes are recognized for preserving chromosomal regions that might otherwise degrade or disappear due to recombination suppression (Lachange, 1967). This preservation is attributed to their ability to retain kinetic activity along the whole chromosomes thus withstanding fragmentation. Notably, it was demonstrated that both in butterflies (which also exhibit a ZW sex-determination system) and in nematodes (which share holocentric chromosomes), inverted meiosis can drive the evolution of new karyotypes (Lukhtanov et al. 2018). Oribatid mites are suggested to exhibit inverted meiosis (Wrensch et al. 1994), thus if these rearrangements happened recently, the sex chromosomes in *H. gibba* might not have had enough time to degenerate.

In our study, the presence of distinct hard edges in the allele frequency plot suggests a singular event, most likely an inversion. Considering the significant role of inversions in contributing to recombination suppression and the formation of sex chromosomes, they align with two possible models i) the sex antagonistic selection model and ii) the deleterious mutation load model.

Sex antagonistic selection is known to be the main driver of recombination suppression in sex-chromosome evolution (Wright et al. 2016). However, a recently proposed model suggests that this might be an overrepresentation (Ironside 2010; Jay et al. 2022). Besides sex antagonistic selection, alternative hypotheses deal with the effect of meiotic drive and genetic drift (Svedberg et al. 2018). However, oribatid mites are known to exhibit large and stable population sizes (Maraun and Scheu 2000) and testing for meiotic drive is difficult (Ponnikas et al. 2018). Nonetheless, insights from comparative genomic analyses in fungi challenge the notion that inversions primarily cause halted recombination on sex chromosomes (Bergero and Charlesworth 2009). Instead, these analyses suggest that recombination suppression was the ancestral state, with inversions emerging as a consequential, rather than primary, factor in recombination suppression. Many forces drive recombination suppression, and identifying a single event is difficult since interactions of different circumstances can result in a loss of recombination forming heterozygous sex chromosomes (Charlesworth et al. 2005; Bergero and Charlesworth 2009; Ponnikas et al. 2018).

In summary, our analysis indicated the absence of highly differentiated regions through coverage analysis. Allele frequencies identified distinct patterns indicative of chromosomal rearrangements, possibly inversions. We hypothesized a relatively recent loss of recombination. Nevertheless, how and why the sex chromosomes might not have differentiated is a complex process influenced by various genetic and structural factors. To obtain a more profound understanding of sex chromosome evolution in mites, it is imperative to expand our investigations to include additional mite species for comprehensive data collection. Considering Acariformes’ position as a basal group within Chelicerata, where diverse mechanisms are observed, our research contributes to enhancing clarity on sex-determination system evolution in basal groups.

## Materials & Methods

Since there is no apparent external dimorphism in these mites, sex was determined by examination of the genitalia as previously described in the literature (Palmer and Norton 1991). The reference genome was from a single individual, and reads were generated from sex-specific pools to perform coverage analysis and allele frequency calculation to determine genomic differences between the sexes.

### Sampling

Animals were collected from ground litter in a coniferous forest in Dahlem, Germany (50.389204 N, 6.568780 E) in April 2021. Specimens were isolated using heat gradient extraction (Kempson, Lloyd, and Ghelardi 1963). *Hermannia gibba* was morphologically identified after Weigmann (Weigmann 2006) and genetically confirmed by cytochrome oxidase I (CO1) sequencing.

### Sample preparation

To minimize contamination from gut contents, mites were starved for at least one week before DNA extraction. Mites were cleansed with a brush in sterile water, in a solution of detergent/water (1:20) (fit GmbH, Zittau, Germany), 70% ethanol (EtOH), and 0.05% bleach (DonKlorix; CP GABA GmbH, Hamburg, Germany), and then rinsed with sterile water. To prevent DNA degradation, mites were stored at -80°C prior to sexing.

### Identification of individuals sexes

Following standard practice for morphological sexing in oribatid mites, the presence of an ovipositor is used for identification of females; spermatophore and missing ovipositor were used to identify male specimens (Figure S3). To perform morphological sexing, the mites were first quickly frozen and then placed onto a microscope slide in 5 µl cooled TNES buffer (400 mmol NaCl, 50 mmol Tris pH 8, 0.5% SDS, 20 mmol EDTA). To expose the ovipositor, genital plates were removed with a dissecting needle, and specimens were dissected completely. To ensure no misidentification and therefore bias to pool data, only clearly identified specimens were used. Sexed individuals were transferred to 190 µl of TNES and frozen at -80°C until *g*DNA extraction. In total 155 specimens were examined with 72 females (∼ 46%) and 83 males (∼ 54%). 30 individuals were pooled to an equimolar amount and sequenced at 3-4X coverage per individual (see Supplementary Table 1). We obtained a total of 38 Gigabases (Gb) of raw data for each pool, with an average of 10.19*10^6^ reads for the female pool and 9.97*10^6^ reads for the male pool.

### gDNA extraction (for extra low input)

We followed the high-molecular-weight *g*DNA extraction protocol for oribatid mites with chitinase digestion (Öztoprak & Bast 2023). In short: chitinase was added to each individual and incubated at 37°C for 60 min. Then Proteinase K was added followed by t yeast tRNA (Invitrogen) and 5M NaCl and 96% ethanol. The solution was incubated at -20°C for 1h or overnight. DNA purification was conducted by washing the pellet twice with fresh and chilled 70% ethanol and eluted in TE buffer. Samples were left to homogenize at 4°C overnight. RNase digestion was conducted and DNA concentration was measured using Qubit Fluorometer v. 4 with the Qubit dsDNA HS Assay kit (ThermoFisher Scientific, Waltham, MA). For pool-sequencing DNA of n = 30 individuals of each sex are pooled to an equimolar amount.

### Long-read sequencing

Single-individual high-molecular-weight DNA was sequenced at Genomics & Transcriptomics Lab (GTL) in Düsseldorf with an ultra-low input protocol for long-read sequencing. Sequencing yielded in total 45.8 Gb of Pacific Biosciences Circular Consensus HiFi reads (Wenger et al. 2019) with an N50 of 15 kilobases (kb).

### Hi-C sequencing

Thirty whole mites were crosslinked in 3% formaldehyde for 1 hour at room temperature with low agitation, followed by quenching in 250 mM glycine for 30 minutes at room temperature with low agitation. The Hi-C library was prepared using the Arima Hi-C+ kit (Arima Genomics); after the addition of lysis buffer, the sample was frozen in liquid nitrogen to crush the mites with a pestle, and the protocol was strictly followed thereafter. The library was sequenced on an Illumina NovaSeq 6000 at the Cologne Center for Genomics (CCG; Cologne, Germany), which generated 88 million pairs of 150-bp reads.

### RNA sequencing

A Direct-zolTM RNA MicroPrep kit (Zymo Research, Irvine, CA) was used to extract total RNA from 10 adults using TRIzol reagent that had been DNase I-treated. With the use of TruSeq Stranded Total RNA with Ribo-Zero Globin, the CCG built an RNA-seq library and sequenced 35x10^6^ pairs of 100-bp reads.

### Pool sequencing

We used n = 30 individuals from each sex for pool sequencing. To meet sample requirements of 3-5-fold coverage for Illumina NovaSeq 6000 Genomic DNA (PCR free -350bp) paired-end DNA sequencing, concentrations were normalized for each individual (Table S1). Genome sequencing and associated library preparation were performed by Novogene Europe (Cambridge, United Kingdom).

### Preliminary analyses of HiFi reads

*k*-mer spectra of the PacBio HiFi reads were built using KAT v2.4.2 (Mapleson et al. 2016) with the modules kat hist and kat gcp (default parameters, k=27). Ploidy was estimated using KMC v3.2.1 (Kokot et al. 2017) with parameters -k 27 -ci 1 -cs 10000 and Smudgeplot v0.2.5 (Ranallo-Benavidez et al. 2020) with default parameters.

### Genome assembly and scaffolding

PacBio HiFi reads were assembled using hifiasm v0.16 (Cheng et al. 2021) with parameter -l 2 Primary contigs were purged of artifactual duplications using minimap2 v2.24 (Li 2021) with parameter -x map-hifi and purge_dups v1.2.5 (Guan et al. 2020) with thresholds -l 6 -m 89 -u 327. Hi-C reads were first trimmed using cutadapt v1.15 (Martin 2011) with parameters -m 10 -a AGATCGGAAGAG -A AGATCGGAAGAG and then pre-processed using BWA v0.7.17 (Li 2013) and hicstuff v3.1.1 (Matthey-Doret et al. 2020) with parameters -e DpnII,HinfI -m iterative -a bwa. The resulting Hi-C sparse matrix was used to scaffold the assembly using instaGRAAL (Baudry et al. 2020) on Galaxy (Galaxy Version 0.1.6; (Galaxy Community 2022) with parameters -l 5 -n 50 -c 1 -N 5. These scaffolds were curated using instaGRAAL-polish with parameters -m polishing -j NNNNNNNNNN. Gaps were filled using TGS-GapCloser v1.1.1 (Xu et al. 2020) and the non-CCS PacBio reads with parameters –tgstype pb –ne. The scaffolds were finally polished by mapping the PacBio HiFi reads using minimap2 v2.24 with parameter -x map-hifi and running HyPo v1.0.3 (Kundu et al. 2019). Assembly was curated using hicstuff for iterative mapping, PretextMap v0.1.9 and PretextView v0.2.5.

### Assembly evaluation

Ortholog completeness was assessed using the tool Benchmarking Universal Single-Copy Orthologs (BUSCO) v.5.0.0 (Manni et al. 2021) against the Arthropoda odb10 lineage (1,066 orthologs) and the Arachnida odb10 lineage (2,934 orthologs). *k*-mer completeness was evaluated using KAT v2.4.2 (Mapleson et al. 2016) and the module kat comp with default parameters. Hi-C reads were mapped to the assembly using BWA (Li 2013) and hicstuff (Matthey-Doret et al. 2020) as previously described. The contact map was generated using the module hicstuff view with the parameter -b 500. The mapped PacBio HiFi reads were mapped to the final scaffolds using minimap2 v2.24 (Li 2021) with parameters -ax map-hifi and the mapped reads were sorted with SAMtools v1.11. The final scaffolds were aligned against the nucleotide database using the Basic Local Alignment Search Tool (BLAST) v2.6.0 (Altschul et al. 1990) with parameters -outfmt “6 qseqid staxids bitscore std sscinames scomnames” -max_hsps 1 -evalue 1e-25. The outputs of minimap2, BLAST, and BUSCO (against the Arachnida odb10 lineage) were provided as input to BlobTools2 v2.3.3 (Challis et al. 2020). Sequences identified as bacteria were subsequently removed.

### Repeat annotation

Transposable elements (TE) were annotated with the EDTA pipeline v1.9.4 (Bell et al. 2022). Long terminal repeats (LTR) were predicted by LTRharvest v1.6.1 (Ellinghaus et al. 2008) and LTR_FINDER v1.07 (Xu and Wang 2007) and then filtered using LTR_retriever v2.9.0 (Ou and Jiang 2018). Helitron transposons were identified using HelitronScanner (Xiong et al. 2014), and other repeats were detected using Generic Repeat Finder v1.0 (Shi and Liang 2019) and TIR-learner (Su et al. 2019). The final TE library was produced after further filtering and repeat annotation with RepeatModeler v2.0.1 (Flynn et al. 2020). The output hardmasked assembly was converted into a softmasked assembly. In addition, telomeres were masked after identification using Telomic Identifier v0.2.1 with parameter -l 5.

### Gene prediction and functional annotation

RNA-seq reads were trimmed using cutadapt v1.15 (Martin 2011) with parameters -m 10 -a AGATCGGAAGAG -A AGATCGGAAGAG and assembled into transcripts using Trinity v2.14 (Grabherr et al. 2011) with default parameters. The genome assembly was then annotated using the pipeline Funannotate v1.8.13 using the trimmed RNA-seq reads and the transcriptome assembly as input. First, the module ‘funannotate train’ pre-processed the RNA-seq reads by mapping them using HISAT2 v2.2.1 (Kim et al. 2019) and providing the output to StringTie v.2.2.1 (Shumate et al. 2022), and processed the transcripts through PASA v2.5.2 (Haas et al. 2003). The initial predictions were provided as input to ‘funannotate predict’ which combined them with predictions from Augustus v3.3.3 (Stanke et al. 2008) using EVidenceModeler v1.1.1 (Haas et al. 2008).

## Data preprocessing

Detailed scripts can be found on Github at https://github.com/TheBastLab/sexdetermination. Raw reads were first checked for quality with FastQC (http://www.bioinformatics.babraham.ac.uk/projects/). Specific adapter sequences of paired-end reads were provided (-a AGATCGGAAGAGCGTCGTGTAGGGAAAGAGTGTAGATCTCGGTGGTCGCCGTATCATT \-a2 GATCGGAAGAGCACACGTCTGAACTCCAGTCACGGATGACTATCTCGTATGCCGTCTTCTGCTTG) and removed and a quality phred score cut off of 30 was performed with TrimGalore v0.6.5 (Krueger et al. 2023). For the allele we used the unmasked reference genome. Trimmed reads were mapped against both versions of the in-house reference genome using BWA v0.7.17 (Li 2013). Aligning sequence reads, clone sequences and assembly contigs with BWA-MEM (Li 2013). Using SAMtools v1.11 (Danecek et al. 2021) SAM files were converted into BAM files and sorted using GATK SortSam v4.1.9.0 (McKenna et al. 2010). Duplicates were then marked using Picard (http://broadinstitute.github.io/picard/) and reindexed.

### Coverage analysis

We used deepTools bamCoverage v3.5.1 (Ramírez et al. 2014) to calculate the coverage as the number of reads per bin. We set a window size of 1 kb and performed the normalization according to the scaling factor *reads per kilobase transcript per million mapped reads* (RPKM; García et al., 2016). The results from the two pools were then merged for further analysis.

### Allele frequency calculation

Allele frequencies (AF) were calculated using MAPGD -pool (Lynch et al. 2014; Ackerman et al. 2017), which uses a maximum likelihood approach to estimate allele frequencies from pooled population genomic data. Probabilities of polymorphism are calculated using the log-likelihood ratio test (LLR) by testing the null hypothesis of no polymorphism. The LLR threshold for identifying a polymorphism corresponds to a significant level of p = 0.00001 (-a 22) (Lynch et al. 2014; Guirao-Rico and González 2021). Allele frequencies were calculated for 4,663,024 million sites, with frequencies calculated based on the major allele in the female and male pools. According to common practice, loci are assumed to be monomorphic/fixed for AF in the range of 0.95 to 1 (Chakraborty et al. 1980).

### Restriction enzyme digestion

Restriction sites were identified using the online tool “GenScript Restriction Enzyme Map Analysis Tools”. (https://www.genscript.com/tools/restriction-enzyme-map-analysis, last accessed on January 21, 2023). Restriction enzyme HindIII was selected for chromosome 1, as the site is exclusive to the female sex. Primers were designed using primer3 with default values (forward: ‘ACACGACAACGCGTCTTTAAT’, reverse: ‘AAAACTCCTGGTTCGCAGTTT’). For PCR we used 1 µl DNA, 10 µl DreamTaq Green PCR Master Mix (2X) (ThermoFisher), 2 µl forward, 2 µl reverse primer and 5 µl ddH_2_O. Each sample was PCR-amplified in a thermal cycler (BentoLab) with initial denaturation at 95°C, followed by 30 cycles of 30 s at 95 °C, 30 s at 60 °C and 1 min at 72 °C. Following ThermoFisher instructions for DNA digestion with a single restriction enzyme, we used 1 µl PCR product (∼2-4 ng/µl), 2 µl 10x Puffer R, 1 µl HindIII, 16 µl ddH_2_O. The reaction was incubated at 37°C for 1.5 hours. The samples were run on a 1.5% agarose gel with 12 µl SYBR™ Green I (ThermoFisher) and TBE 1×.

## Supplementary Information

### Supplementary Figures

**Figure S1.**
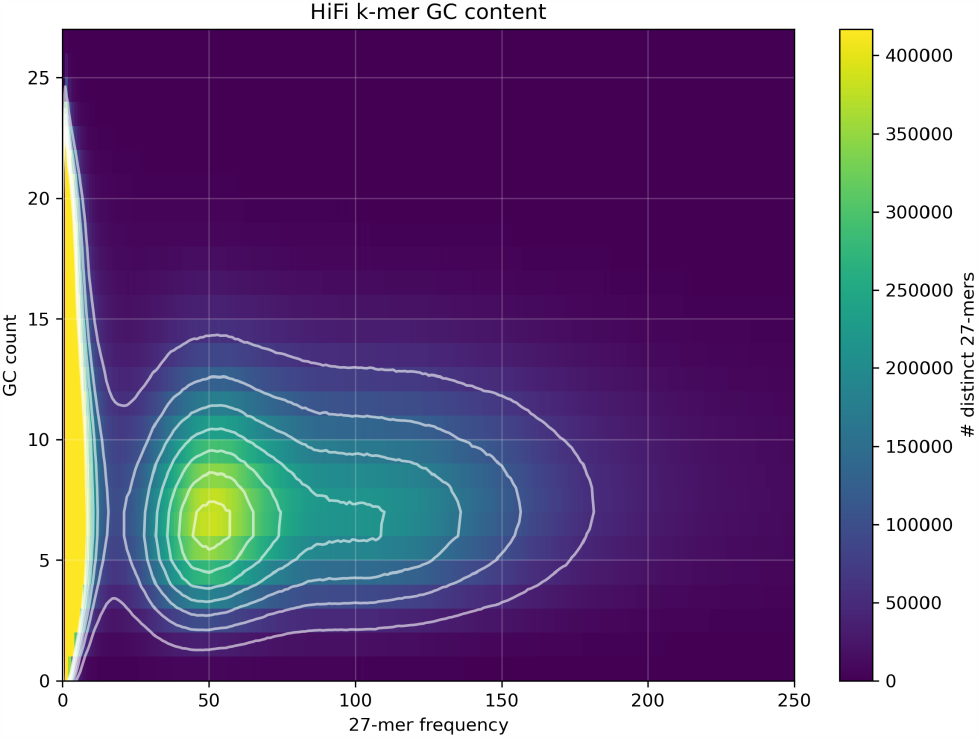
Analysis of GC content and frequency of 27-mers. There are two peaks at frequencies of 50 and 100 (respectively, heterozygous and homozygous peaks) with a GC content mainly between 4 and 9. This plot shows no bacterial contaminants.

**Figure S2.**
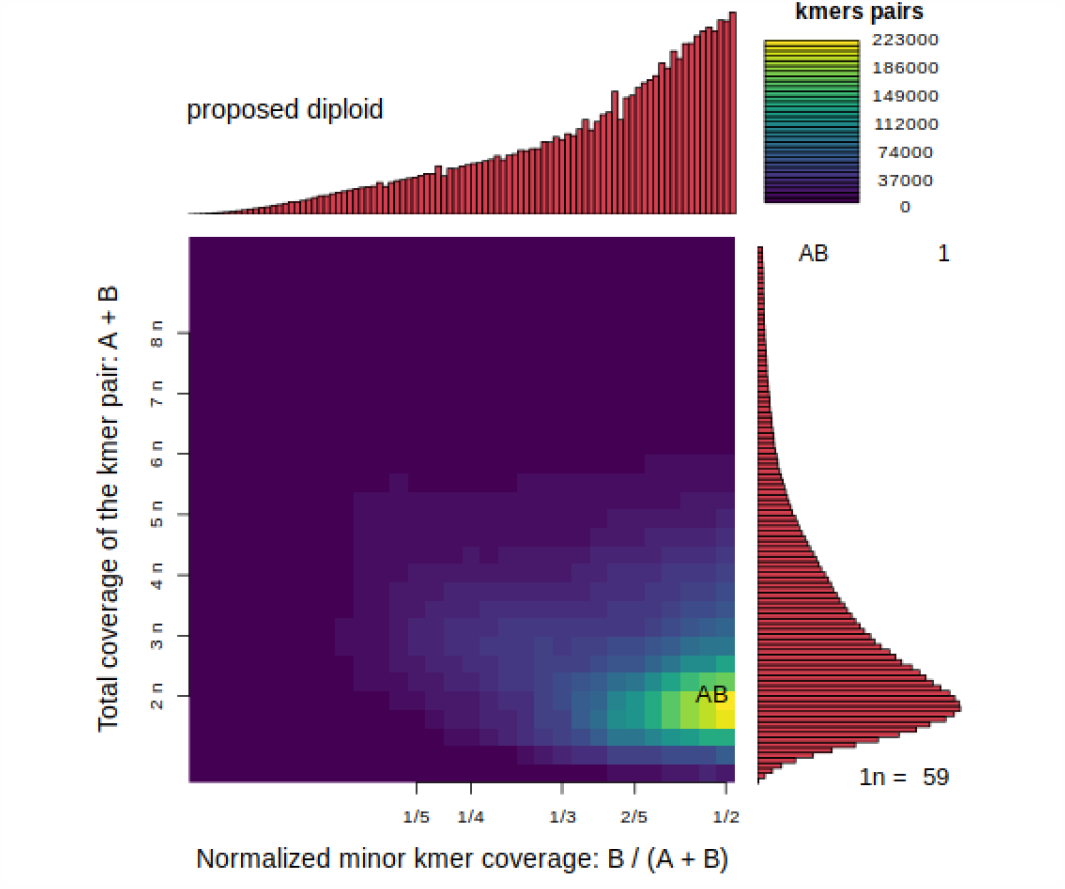
*k*-mer analysis of ploidy. The strong AB smudge predicts that the species is diploid.

**Figure S3.**
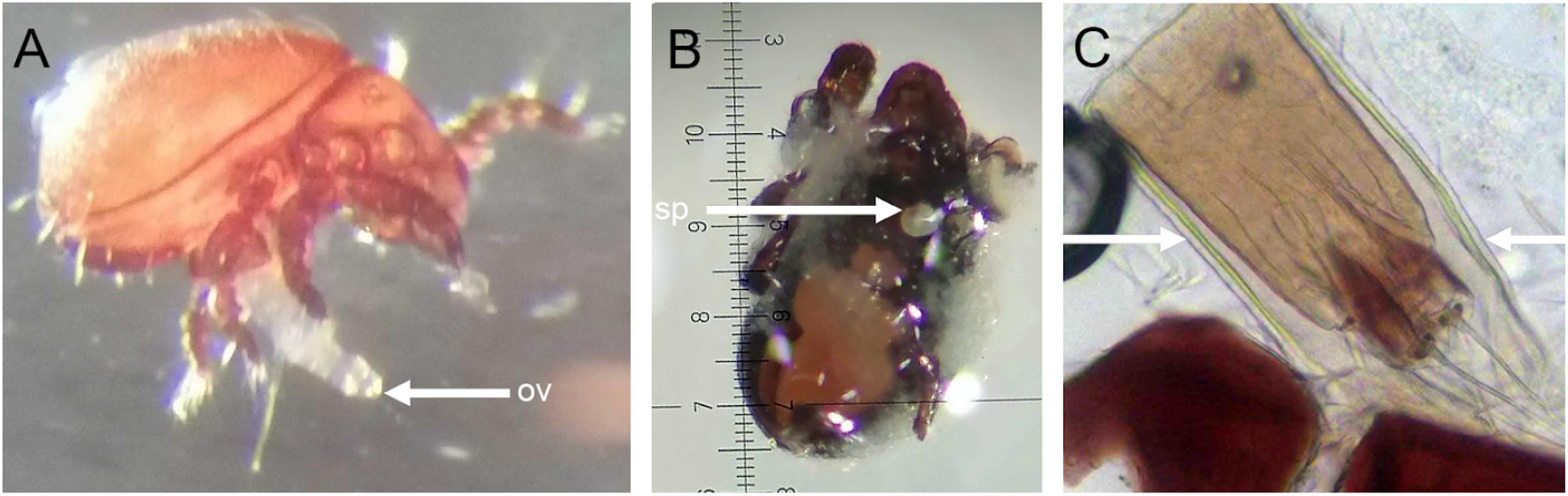
Overview of morphological sex determination in *Hermannia gibba* (A) Arrow points to the ovipositor (ov) of an adult female (B) Arrows shows spermatophore (x5 magnification) (sp) of an adult male (C) In between arrows dissected ovipositor.

**Figure S4.**
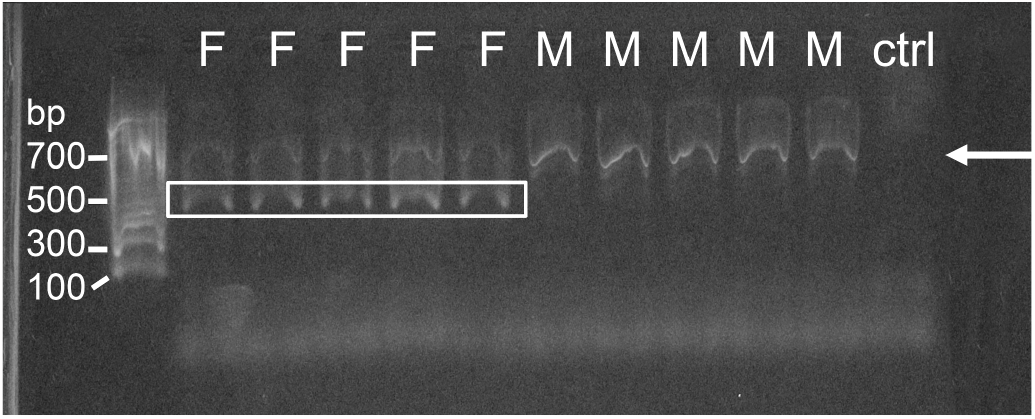
Restriction enzyme digestion approach was conducted using following individuals with corresponding cDNA concentrations: female ID: 10: 6.20 ng/µl, 111: 2.85 ng/µl, 74: 2.79 ng/µl, 75: 6.06 ng/µl, 77: 3.95 ng/µl; male ID: 65: 5.25ng/µl, 67: 3.56 ng/µl, 68: 4.08 ng/µl, 70: 3.06ng/µl, 81: 5.39 ng/µl). The experiment targeted a female-specific SNP in the Sex Determining Region (SDR) on chromosome 1 using the HindIII restriction enzyme. In both the male and female samples, PCR amplification resulted in a 710-base pair product. After digestion, only female DNA produced two fragments (380 and 330 base pairs), confirming the presence of a female-heterogametic ZW sex-determination system in *H. gibba*.

### Restriction enzyme digestion for sex verification

To additionally verify the identification of a ZW system and to be able to sex individuals collected from natural populations without the need for time-consuming morphological analyses, we designed a restriction enzyme digestion assay for *Hermannia gibba*. Therefore, we screened for the presence of a female-specific SNP in the SDR on chromosome 1, which is located within a restriction site of the HindIII restriction enzyme. PCR amplification of both male and female products has an expected size of 710 base pairs. After restriction enzyme digestion, the PCR product was cleaved as expected into two fragments with sizes of 380 and 330 base pairs, in the female DNA pool only (Figure S4). This confirmed the presence of a female-heterogametic ZW sex-determination system in *H. gibba*.

## Supplementary Graphs and Tables

**Table S1.** Morphological sex determination of 30 specimens of both sexes, DNA concentration per individual (ng/µl), pooling amount per individual for equimolar DNA concentrations for each pool (µl)

| No. | Sex | Sample-ID | Qubit (ng/μl) | Pooling amount (μl) | No. | Sex | Sample-ID | Qubit (ng/μl) | Pooling amount (μl) |
| --- | --- | --- | --- | --- | --- | --- | --- | --- | --- |
| 1 | Female | 1 | 4,45 | 7,19 | 1 | Male | 65 | 3,05 | 10,49 |
| 2 | Female | 2 | 5,34 | 5,99 | 2 | Male | 66 | 3,95 | 8,1 |
| 3 | Female | 3 | 5,07 | 6,31 | 3 | Male | 67 | 4,49 | 7,13 |
| 4 | Female | 4 | 7,08 | 4,52 | 4 | Male | 68 | 3,92 | 8,16 |
| 5 | Female | 5 | 5,25 | 6,1 | 5 | Male | 70 | 5,23 | 6,12 |
| 6 | Female | 6 | 6,84 | 4,68 | 6 | Male | 71 | 3,71 | 8,63 |
| 7 | Female | 7 | 3,96 | 8,08 | 7 | Male | 81 | 2,97 | 10,77 |
| 8 | Female | 8 | 4,71 | 6,79 | 8 | Male | 86 | 3,34 | 9,58 |
| 9 | Female | 9 | 5,51 | 5,81 | 9 | Male | 87 | 3,61 | 8,86 |
| 10 | Female | 10 | 6,2 | 5,16 | 10 | Male | 88 | 3,5 | 9,14 |
| 11 | Female | 74 | 2,79 | 11,47 | 11 | Male | 89 | 2,32 | 13,79 |
| 12 | Female | 75 | 6,06 | 5,28 | 12 | Male | 90 | 4,17 | 7,67 |
| 13 | Female | 77 | 3,95 | 8,1 | 13 | Male | 92 | 3,91 | 8,18 |
| 14 | Female | 78 | 3,12 | 10,26 | 14 | Male | 93 | 3,96 | 8,08 |
| 15 | Female | 105 | 5,67 | 5,64 | 15 | Male | 94 | 3,99 | 8,02 |
| 16 | Female | 110 | 5,19 | 6,17 | 16 | Male | 95 | 2,37 | 13,5 |
| 17 | Female | 111 | 2,85 | 11,23 | 17 | Male | 100 | 6,66 | 4,8 |
| 18 | Female | 114 | 5,92 | 5,41 | 18 | Male | 107 | 6,59 | 4,86 |
| 19 | Female | 125 | 5,49 | 5,83 | 19 | Male | 112 | 3,32 | 9,64 |
| 20 | Female | 126 | 4,48 | 7,14 | 20 | Male | 113 | 2,15 | 14,88 |
| 21 | Female | 127 | 6,02 | 5,32 | 21 | Male | 115 | 2,39 | 13,39 |
| 22 | Female | 128 | 4,99 | 6,41 | 22 | Male | 117 | 4,03 | 7,94 |
| 23 | Female | 129 | 3,66 | 8,74 | 23 | Male | 118 | 7,93 | 4,04 |
| 24 | Female | 137 | 5,3 | 6,04 | 24 | Male | 119 | 6,83 | 4,69 |
| 25 | Female | 138 | 3,86 | 8,29 | 25 | Male | 120 | 5,24 | 6,11 |
| 26 | Female | 139 | 5,38 | 5,95 | 26 | Male | 121 | 3,28 | 9,76 |
| 27 | Female | 140 | 2,82 | 11,35 | 27 | Male | 122 | 4,97 | 6,44 |
| 28 | Female | 142 | 6,16 | 5,19 | 28 | Male | 123 | 4,52 | 7,08 |
| 29 | Female | 143 | 4,18 | 7,66 | 29 | Male | 124 | 6,77 | 4,73 |
| 30 | Female | 144 | 7,15 | 4,48 | 30 | Male | 130 | 4,67 | 6,85 |

## Author contributions

Conceptualization: JB

Resources: HÖ, SW

Methodology: JB, HÖ, SW, NG, DLJ

Investigation: SW, HÖ

Visualization: SW, NG, HÖ

Funding acquisition: JB

Project administration: JB, DLJ, HÖ

Supervision: JB, DLJ, HÖ, NG

Data curation: NG

Formal Analyses: SW, NG

Writing – original draft: SW, HÖ, with input from all authors

Writing – review & editing: SW, JB

## Data and materials availability

All code for the analyses can be found at https://github.com/TheBastLab/sexdetermination. The genome, together with its annotation is available at X

